# metamicrobiomeR: an R package for analysis of microbiome relative abundance data using zero-inflated beta GAMLSS and meta-analysis across studies using random effect models

**DOI:** 10.1101/294678

**Authors:** Nhan Thi Ho, Fan Li

## Abstract

**Background:** The rapid growth of high-throughput sequencing-based microbiome profiling has yielded tremendous insights into human health and physiology. Data generated from high-throughput sequencing of 16S rRNA gene amplicons are often preprocessed into composition or relative abundance. However, reproducibility has been lacking due to the myriad of different experimental and computational approaches taken in these studies. Microbiome studies may report varying results on the same topic, therefore, meta-analyses examining different microbiome studies to provide robust results are important. So far, there is still a lack of implemented methods to properly examine differential relative abundances of microbial taxonomies and to perform meta-analysis examining the heterogeneity and overall effects across microbiome studies.

**Results:** We developed an R package ‘*metamicrobiomeR’* that applies Generalized Additive Models for Location, Scale and Shape (GAMLSS) with a zero-inflated beta (BEZI) family (GAMLSS-BEZI) for analysis of microbiome relative abundance datasets. Both simulation studies and application to real microbiome data demonstrate that GAMLSS-BEZI well performs in testing differential relative abundances of microbial taxonomies. Importantly, the estimates from GAMLSS-BEZI are log(odds ratio) of relative abundances between groups and thus are comparable between microbiome studies. As such, we also apply random effects meta-analysis models to pool estimates and their standard errors across microbiome studies. We demonstrate the meta-analysis workflow and highlight the utility of our package on four studies comparing gut microbiomes between male and female infants in the first six months of life.

**Conclusions:** GAMLSS-BEZI allows proper examination of microbiome relative abundance data. Random effects meta-analysis models can be directly applied to pool comparable estimates and their standard errors to evaluate the heterogeneity and overall effects across microbiome studies. The examples and workflow using our *metamicrobiomeR* package are reproducible and applicable for the analyses and meta-analyses of other microbiome studies.

## BACKGROUND

The rapid growth of high-throughput sequencing-based microbiome profiling has yielded tremendous insights into human health and physiology. However, interpretation of microbiome studies have been hampered by a lack of reproducibility in part due to the variety of different study designs, experimental approaches, and computational methods used [1, 2]. Microbiome studies may report varying results on the same topic. Therefore, meta-analyses examining different microbiome studies are critical to provide robust results. Meta-analysis studies pooling individual sample data across studies for pooled analysis of all samples or processing of all samples together followed by analysis of each study separately have revealed some consistent microbial signatures in certain conditions such as inflammatory bowel disease (IBD) and obesity [3–9]. Software has been developed for the analysis and meta-analysis of microbiome data [10]. However, these studies do not explicitly model mirobiome relative abundance data using an appropriate statistical method and do not examined between-group-comparison overall pooled effects in the meta-analysis. For example, a recently published meta-analysis study examined differential relative abundances of bacterial taxonomies of each study separately using Wilcoxon’s tests but did not examine overall pooled effects across studies [8]. Moreover, meta-analysis pooling estimates across studies is not practical if studies utilize some methods such as Wilcoxon’s tests or Random Forest (RF) as there are no test statistics or estimates that are applicable for pooling.

Data generated from high-throughput sequencing of 16S rRNA gene amplicons are often preprocessed into relative abundance. Microbiome relative abundances are compositional data which range from zero to one and are generally zero-inflated. To test for differences in relative abundance of microbial taxonomies between groups, methods such as bootstrapped non-parametric t-tests or Wilcoxon tests (not suitable for longitudinal data and covariate adjustment) [11–13] and linear or linear mixed effect models (LM) [14, 15] (suitable for longitudinal data and covariate adjustment) have been widely used. However, these methods do not address the actual distribution of the microbiome relative abundance data, which resemble a zero-inflated beta distribution. Transformations (e.g. arcsin square root) of relative abundance data to make it resemble continuous data to use in LM has been proposed by Morgan et al (implemented in MaAsLin software) [16] and has been widely used to test for differential relative abundances [17–20]. However, this adjustment does not address the inflation of zero values in microbiome relative abundance data. More recently, methods adapted from the RNA-seq field that account for zero inflation and utilize Poisson or negative binomial models have shown some promise in differential abundance testing of microbiome datasets [21, 22]. However, these methods are applicable to absolute abundance, which are count data and largely vary between studies due to the difference in experimental and computational approaches making the estimates from these methods incomparable across studies. Therefore, standard meta-analysis approaches may not be directly applicable to pool estimates and their standard errors from these methods across studies.

Here, we developed an R package “*metamicrobiomeR”* that applies Generalized Additive Models for Location, Scale and Shape (GAMLSS) [23] with a zero-inflated beta (BEZI) family (GAMLSS-BEZI) for the analysis of microbial taxonomy relative abundance data. With BEZI family, this model allows direct and proper examination of microbiome relative abundance data, which resemble a zero-inflated beta distribution. This model also allows adjustment for covariates and can be used for both non-longitudinal and longitudinal studies. We evaluated the performance of GAMLSS-BEZI using simulation studies and real microbiome data. Importantly, the estimates from GAMLSS-BEZI are log(odds ratio) of relative abundances between groups and thus are comparable across microbiome studies and can be directly combined using standard meta-analysis approaches. As such, we apply random effect meta-analysis models to pool the estimates and standard errors as part of the “*metamicrobiomeR”* package. This approach allows examination of study-specific effects, heterogeneity between studies, and the overall pooled effects across studies. Finally, we provide examples and sample workflows for both components of the “*metamicrobiomeR”* package. Specifically, we use GAMLSS-BEZI to compare relative abundances of the gut microbial taxonomies of male versus female infants ≤ 6 months of age while adjusting for feeding status and infant age at time of sample collection and demonstrate the application of the random effect meta-analysis component on four studies of the infant gut microbiome.

## IMPLEMENTATION

### GAMLSS-BEZI for the analysis of bacterial taxa relative abundance and bacterial functional pathway relative abundance data

Relative abundances of bacterial taxa at various taxonomic levels (from phylum to genus or species) are obtained via the “*summarize_taxa.py”* script in QIIME1 [13]. Bacterial functional pathway abundances (e.g. Kyoto Encyclopedia of Genes and Genomes (KEGG) pathway level 1 to 3) are obtained from metagenome prediction analysis using PICRUSt [24]. In the *taxa.compare* function, all bacterial taxa or pathway data are first filtered to retain features with mean relative abundance ≥ relative abundance threshold (e.g. ≥0.005%) and with prevalence ≥ prevalence threshold (e.g. present in ≥ 5% of the total number of samples). This pre-filtering step has been shown to improve performance of various differential abundance detection strategies [25]. A filtered data matrix is then modeled by GAMLSS-BEZI and (mu) logit link and other default options using the R package gamlss version 5.0-5 [23]. A subject random effect can be added to the model for longitudinal data. For performance evaluation, LM and LM with arcsin squareroot transformation (LMAS) were also implemented in the function *taxa.compare*.

Multiple testing adjustment can be done using different methods (False Discovery Rate (FDR) control by default). Below is an example call of the *taxa.compare* function: *taxa.compare(taxtab=taxtab, propmed.rel=‘gamlss’,transform=‘none’, comvar=‘gender’,adjustvar=c(‘age.sample’,‘feeding’),longitudinal=‘yes’, percent.filter=0.05,relabund.filter=0.00005, p.adjust.method=‘fdr’)*

For visualization purposes and subsequent meta-analysis, the output from *taxa.compare* comprises matrices containing coefficients, standard errors, p-values and multiple testing adjusted p-values of all covariates in the models for each bacterial taxon or pathway.

### Meta-analysis across studies using random effects models

The adjusted estimates for bacterial taxa or pathway relative abundances from the GAMLSS models are log(odds ratio) of relative abundances between groups and thus are comparable across microbiome studies. Therefore, standard meta-analysis approaches can be directly applied. In the *meta.taxa* function, random effects meta-analysis models pooling adjusted estimates and standard errors with inverse variance weighting and the DerSimonian–Laird estimator for between-study variance are implemented to estimate the overall effects, corresponding 95% confidence intervals (95% CI) and heterogeneity across studies. A fixed effect meta-analysis model is also implemented for comparison. Meta-analyses is performed only for taxa or pathways observed in ≥ a specified percentage threshold (e.g. 50%) of the total number of included studies. An example call to *meta.taxa* using the output data matrices combined from multiple calls to the *taxa.compare* function is shown below: *meta.taxa(taxcomdat=combined.taxa.compare.output,summary.measure=‘RR’,pool.var=‘id’,st udylab=‘study’,backtransform=FALSE,percent.meta=0.5,p.adjust.method=‘fdr’)*

The output from *meta.taxa* consists of pooled estimates, standard errors, 95% CI, pooled p-values and multiple testing adjusted pooled p-values of all covariates for each bacterial taxon or pathway.

### Additional microbiome measures

Random Forest (RF) modeling of gut microbiota maturity has been widely used to characterize development of the microbiome over chronological time [12, 14, 26]. Adapting from the original approach of Subramanian et al [14], in the *microbiomeage* function, relative abundances of bacterial genera that were detected in the Bangladesh data [14] and in the data of other studies to be included were regressed against infant chronological age using a RF model on a predefined training dataset of the Bangladesh study. This predefined training set includes 249 samples collected monthly from birth to 2 years of age from 11 Bangladeshi healthy singleton infants.

The RF training model fit based on relative abundances of these shared bacterial genera was then used to predict infant age on the test data of the Bangladesh study and the data of each other study to be included. The predicted infant age based on relative abundances of these shared bacterial genera in each study is referred to as gut microbiota age (Additional file 1).

All implemented functions in the *metamicrobiomeR* package are summarized in Table 1.

**Table 1:** Summary of all implemented functions in the *metamicrobiomeR* package.

| Functions | Description |
| --- | --- |
| *taxa.filter* | Filter relative abundances of bacterial taxa or pathways using prevalence and abundance thresholds |
| *taxa.meansdn* | Summarize mean, standard deviation of abundances and number of subjects by groups for all bacterial taxa or pathways |
| *taxa.mean.plot* | Plot mean abundance by groups (from *taxa.meansdn* output) |
| *taxa.compare* | Compare relative abundances of bacterial taxa at all levels using Generalized Additive Models for Location, Scale and Shape (GAMLSS) with a zero-inflated beta (BEZI) family (GAMLSS-BEZI) or linear/linear mixed effect models (LM) or linear/linear mixed effect models with arcsin squareroot transformation (LMAS) |
| *pathway.compare* | Compare relative abundances of bacterial functional pathways at all levels using GAMLSS-BEZI or LM or LMAS. Compare of log(absolute abundances) of bacterial functional pathways at all levels using LM |
| *taxcomtab.show* | Display the results of relative abundance comparison (from *taxa.compare* or *pathway.compare* outputs) |
| *meta.taxa* | Perform meta-analysis of relative abundance estimates of bacterial taxa or pathways (either from GAMLSS-BEZI or LM or LMAS) across studies (from combined *taxa.compare/pathway.compare* outputs of all included studies) using random effect and fixed effect meta-analysis models |
| *metatab.show* | Display meta-analysis results of bacterial taxa or pathway relative abundances (from *meta.taxa* output) |
| *meta.niceplot* | Produce nice combined heatmap and forest plot for meta-analysis results of bacterial taxa and pathway relative abundances (from *metatab.show* output) |
| *alpha.compare* | Calculate average alpha diversity indexes for a specific rarefaction depth, standardize and compare alpha diversity indexes between groups |
| *microbiomeage* | Predict microbiota age using Random Forest model based on relative abundances of bacterial genera shared with the Bangladesh study [14] |
| *read.multi* | Read multiple files in a path to R |

### RESULTS and DISCUSSION

#### Performance of GAMLSS-BEZI: simulation studies

Simulation studies were performed to evaluate type I error and power of GAMLSS-BEZI for testing differential relative abundances of microbial taxonomies as compared to LM with arcsin squareroot transformation (LMAS) (implemented in MaAsLin software [16]). LMAS was chosen for comparison with GAMLSS-BEZI because it is a commonly used approach for microbiome differential relative abundance testing and similarly to GAMLSS-BEZI, it allows covariate adjustment and can be used for longitudinal or non-longitudinal data. Simulations of zero-inflated relative abundance data were based on the R package “gamlss.dist” version 5.0-3 [27].

#### Type I error

Three scenarios were simulated to mimic typical case-control microbiome studies of small (number of cases (n1) =number of controls (n2)=10), medium (n1=n2=50) and large (n1=n2=200) sample sizes. The same location parameter values (mu1=mu2=0.5), precision parameter values (sigma1=sigma2=5) and parameter values modelling the probability at zero (nu1=nu2=0.5) were used for case and control groups. These parameters were chosen to generate the data that mimics real microbiome relative abundance data used in our later examples. The simulations were repeated 10000 times for each sample size. Type I error was calculated for three different alpha levels of 0.01, 0.05 and 0.1. Under these conditions, type I errors were well controlled by both GAMLSS-BEZI and LMAS (Table 2).

**Table 2.**
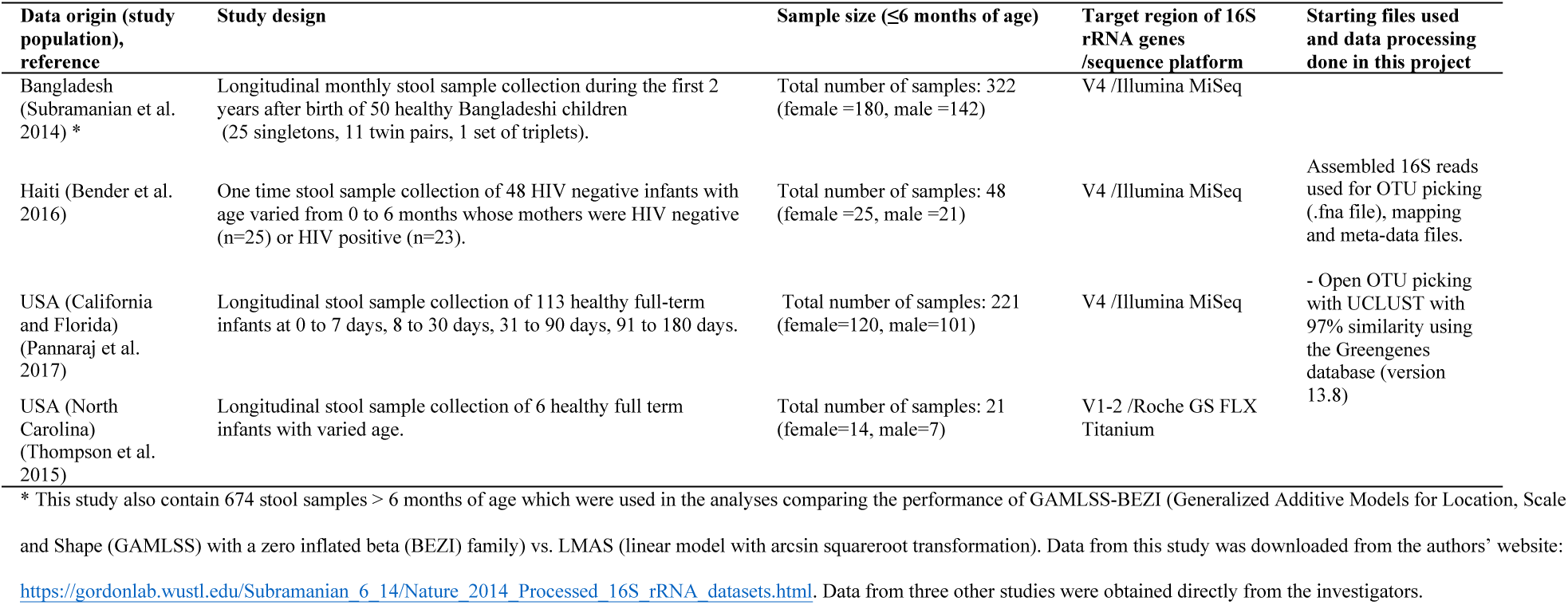
Type I error of GAMLSS-BEZI and LMAS: simulation studies.

The same location parameter values (mu1=mu2=0.5), precision parameter values (sigma1=sigma2=5) and parameter values modelling the probability at zero (nu1=nu2=0.5) were used for case and control groups. Simulations were repeated 10000 times.

GAMLSS-BEZI: Generalized Additive Models for Location, Scale and Shape (GAMLSS) with a zero inflated beta (BEZI) family; LMAS: linear model with arcsin squareroot transformation (implemented in the software MaAsLin).

#### Power

We simulated 5000 bacterial species including 1000 species with no difference between case and control groups (mu1=mu2=0.5) and 4000 species with true difference between case and control groups. Four scenarios of true difference were used (1000 species each): mu1=0.5 vs. mu2=0.4, mu1=0.5 vs. mu2=0.3, mu1=0.5 vs. mu2=0.2, mu1=0.2 vs. mu2=0.1. Other parameters were the same for case and control groups as above. A sample size of n=100 for both case and control groups was used in all scenarios. Performance of GAMLSS-BEZI and LMAS was evaluated based on the receiver operating characteristic (ROC) curve for identifying species with differential abundance between case and control groups. The analysis for the ROC and area under the curve (AUC) was done using the R package pROC version 1.10.0 [27]. GAMLSS-BEZI (AUC=91.1%, 95% CI=(90.2%, 91.9%)) significantly outperformed LMAS (AUC=86.2%, 95% CI=(85.2%, 87.3%)) (DeLong’s test p-value < 2.2e-16) (Figure 1).

**Figure 1.**
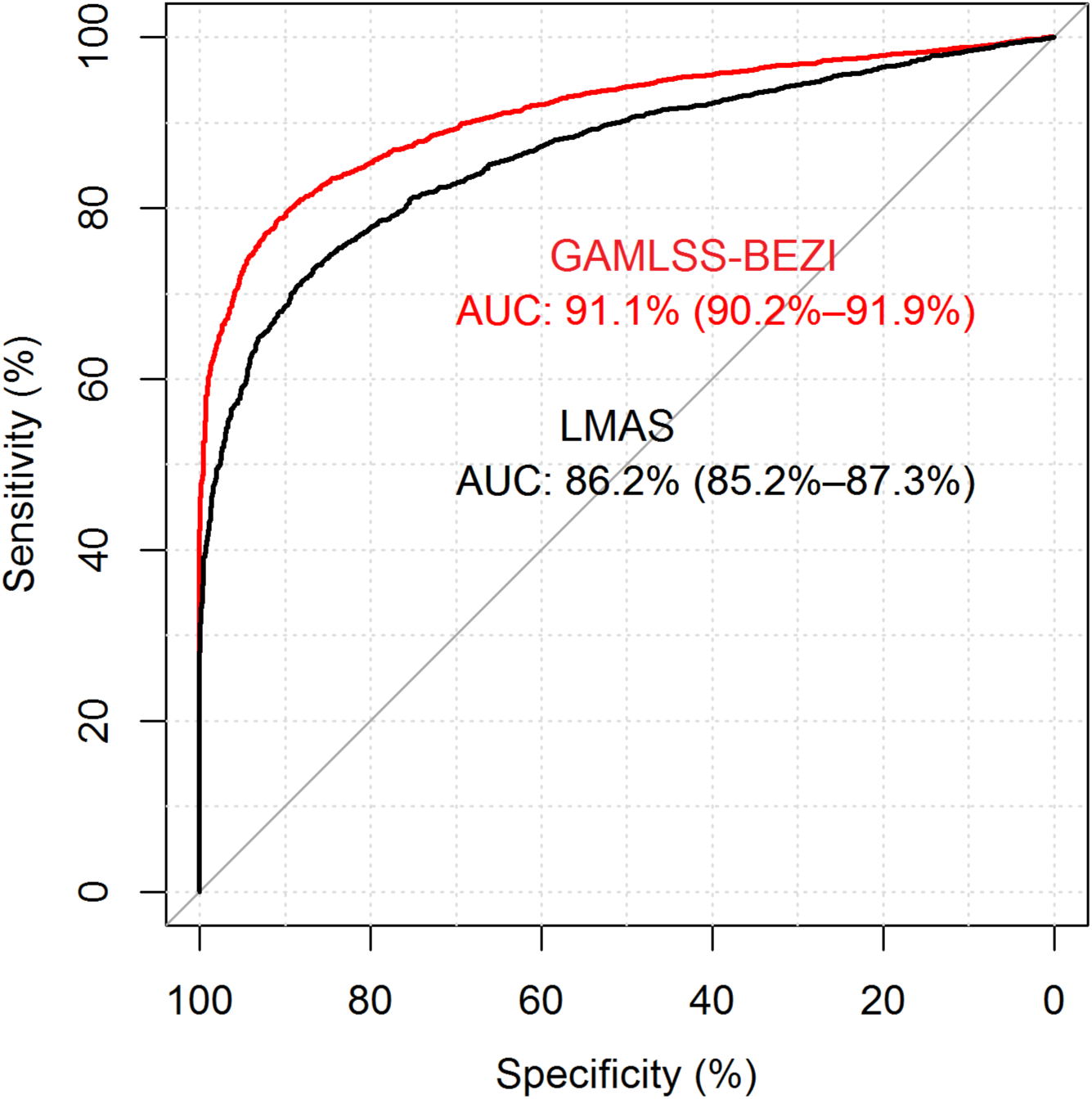
ROC curves for identifying simulated bacterial species with differential relative abundances by GAMLSS-BEZI and LMAS. 5000 bacterial species were simulated including 1000 species with no difference between case and control groups (location parameter values for cases (mu1) = location parameter values for controls (mu2)=0.5) and 4000 species with true difference between case and control groups. Four scenarios for true difference were simulated (1000 species each): mu1=0.5 vs. mu=0.4, mu1=0.5 vs. mu2=0.3, mu1=0.5 vs. mu2=0.2, mu1=0.2 vs. mu2=0.1. Precision parameter values (sigma1=sigma2=5) and parameter values modeling the probability at zero (nu1=nu2=0.5) were the same for case and control groups. A sample size of n=100 for both case and control groups was used in all scenarios. The figure shows ROC curves and AUC with 95% confidence interval. ROC curve: receiver operating characteristic (ROC) curve; AUC: area under the curve; GAMLSS-BEZI: Generalized Additive Models for Location, Scale and Shape (GAMLSS) with a zero inflated beta (BEZI) family; LMAS: linear model with arcsin squareroot transformation (implemented in the software MaAsLin).

#### Performance of GAMLSS-BEZI: application to real microbiome data

Using data from a study of Bangladeshi infants [14], which included 996 stool samples collected monthly from birth to 2 years of life in 50 subjects, we demonstrate that GAMLSS-BEZI out-performs LMAS in detecting differential relative abundances between various grouping variables.

*Example 1: Comparison between non-exclusively breastfed (non-EBF) vs. exclusively breastfed (EBF) infants ≤ 6 months of age*

Figure 2 showed average of relative abundance of bacterial phyla in non-EBF and EBF infants from birth to 6 months of age. A higher abundance of Proteobacteria, Firmicutes, and Bacteroidetes as well as a lower abundance of Actinobacteria was observed in non-EBF versus EBF infants. GAMLSS-BEZI was able to detect a significant difference in all four of these phyla whereas LMAS could only detect a significant difference in three phyla (Table 3).

**Figure 2.**
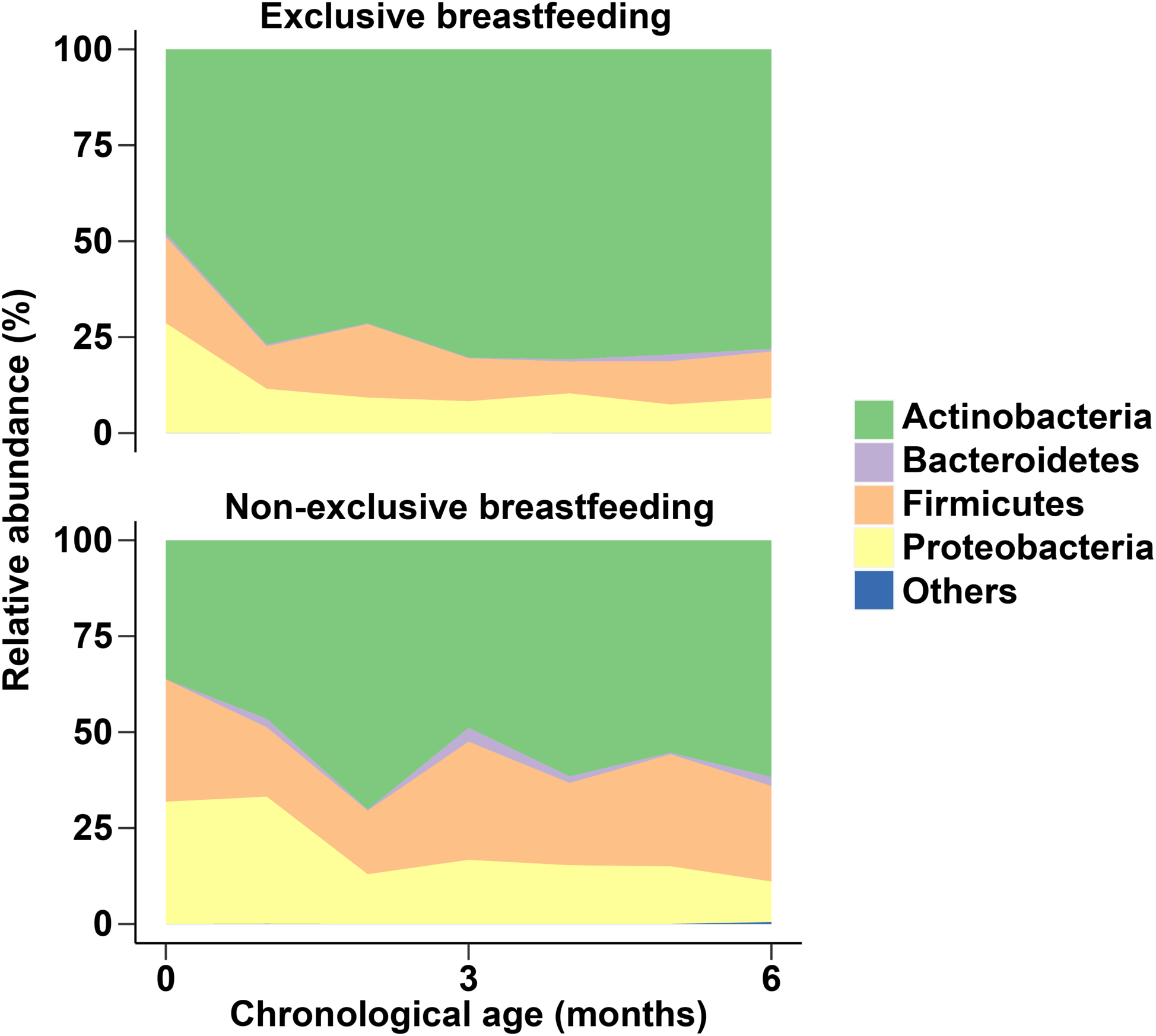
Relative abundances of bacterial phyla in non-exclusively breastfed vs. exclusively breastfed infants ≤ 6 months of age. Data from Bangladesh study.

**Table 3.** Results of GAMLSS-BEZI and LMAS: real microbiome data example 1 (non-EBF vs. EBF).

| GAMLSS-BEZI | LMAS |
| --- | --- |

| Bacterial phyla | Estimate | 95% Lower limit | 95% Upper limit | p-value | FDR adjusted p-value | Estimate | 95% Lower limit | 95% Upper limit | p-value | FDR adjusted p-value |
| --- | --- | --- | --- | --- | --- | --- | --- | --- | --- | --- |
| Actinobacteria | -0.37 | -0.65 | -0.10 | **0.0083** | 0.0166 | -0.13 | -0.23 | -0.03 | **0.0088** | 0.0207 |
| Bacteroidetes | 0.26 | 0.00 | 0.53 | **0.0499** | 0.0499 | 0.03 | 0.00 | 0.05 | **0.0292** | 0.0390 |
| Firmicutes | 0.24 | 0.00 | 0.47 | **0.0468** | 0.0499 | 0.07 | 0.00 | 0.14 | 0.0668 | 0.0668 |
| Proteobacteria | 0.37 | 0.11 | 0.64 | **0.0053** | 0.0166 | 0.10 | 0.02 | 0.17 | **0.0103** | 0.0207 |

Data from Bangladesh study. Comparison of bacterial relative abundances at phylum level between non-exclusively breastfed (non-EBF) vs. exclusively breastfed (EBF) infants ≤ 6 months of age using GAMLSS-BEZI vs. LMAS. Significant p-values (<0.05) are in bold. GAMLSS-BEZI: Generalized Additive Models for Location, Scale and Shape (GAMLSS) with a zero inflated beta (BEZI) family; LMAS: linear model with arcsin squareroot transformation (implemented in the software MaAsLin); FDR: false discovery rate.

*Example 2: Comparison between infants from 6 months to 2 years of age introduced to solid food after 5 months vs. before 5 months*

Figure 3 showed average of relative abundance of bacterial phyla in two groups of in infants from 6 months to 2 years of age who was introduced to solid food after 5 months vs. before 5 months of life. Lower relative abundances of Firmicutes, Bacteroidetes and higher relative abundance of Actinobacteria was observed in infants with solid food introduction after 5 months. GAMLSS-BEZI detected all three of these differences whereas LMEM could only detect a significant difference in one phylum (Table 4).

**Figure 3.**
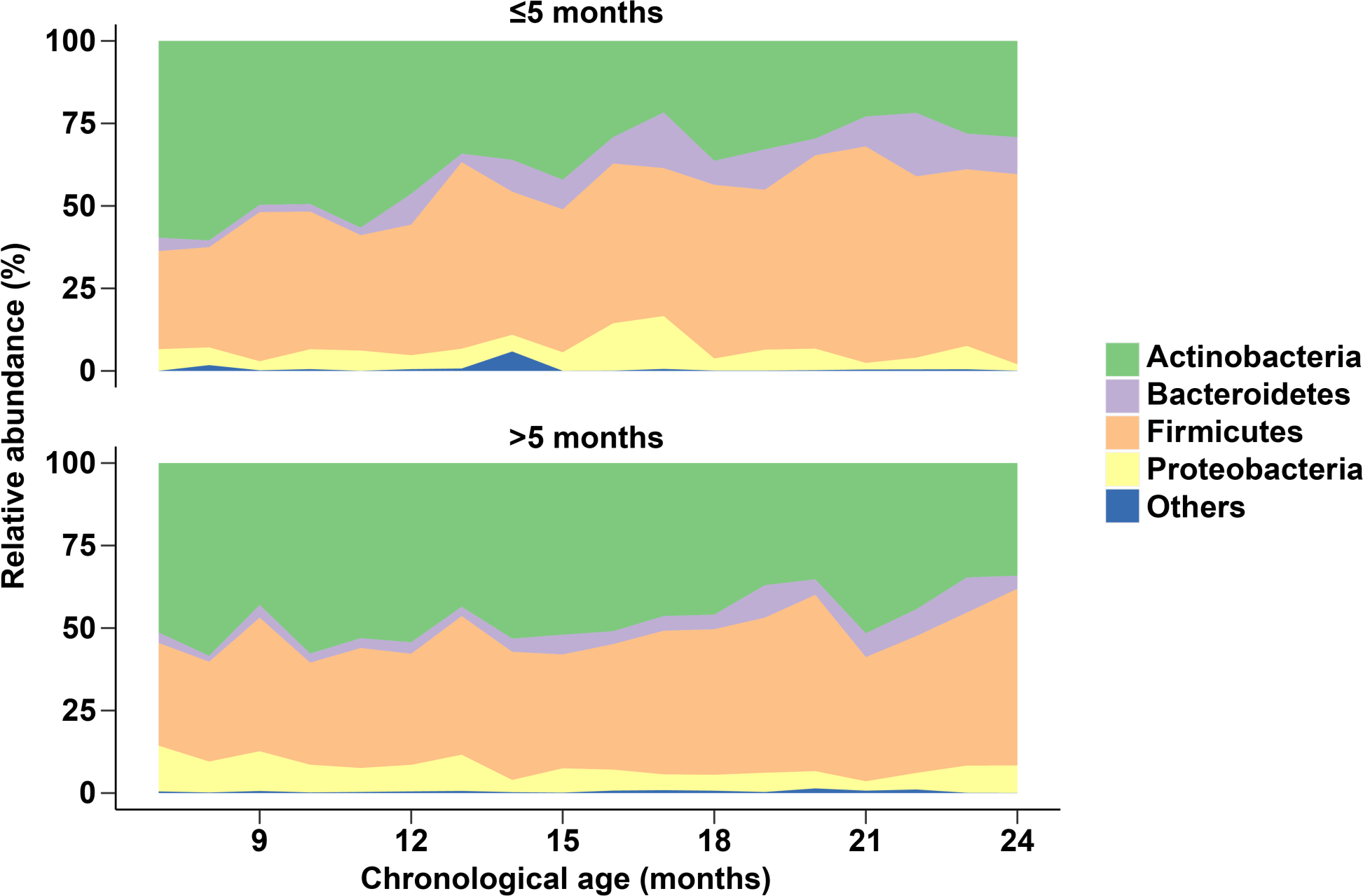
Relative abundances of bacterial phyla in infants from 6 months to 2 years of age with solid food introduction after 5 months vs. before 5 months. Data from Bangladesh study.

Example 1 and 2 demonstrate the increased sensitivity of GAMLSS-BEZI in detecting bacterial taxa with differential relative abundances as compared to LMAS.

**Table 4.** Results of GAMLSS-BEZI and LMAS: real microbiome data example 2 (solid food introduction after 5 months vs. before 5 months of age).

| Bacterial phyla | GAMLSS-BEZI |  |  |  |  | LMAS |  |  |  |  |
| --- | --- | --- | --- | --- | --- | --- | --- | --- | --- | --- |
|  | Estimate | 95% Lower limit | 95% Upper limit | p-value | FDR adjusted p-value | Estimate | 95% Lower limit | 95% Upper limit | p-value | FDR adjusted p-value |
| Actinobacteria | 0.19 | 0.04 | 0.34 | **0.0119** | 0.0208 | 0.05 | -0.06 | 0.16 | 0.3451 | 0.3451 |
| Bacteroidetes | -0.26 | -0.42 | -0.10 | **0.0018** | 0.0070 | -0.05 | -0.09 | -0.01 | **0.027** | 0.1079 |
| Firmicutes | -0.16 | -0.30 | -0.03 | **0.0156** | 0.0208 | -0.04 | -0.12 | 0.04 | 0.3168 | 0.3451 |
| Proteobacteria | 0.14 | -0.02 | 0.30 | 0.0861 | 0.0861 | 0.02 | -0.02 | 0.07 | 0.2916 | 0.3451 |

Data from Bangladesh study. Comparison of bacterial relative abundances at phylum level between infants from 6 months to 2 years of age with solid food introduction after 5 months vs. before 5 months of age using GAMLSS-BEZI vs. LMAS. Significant p-values (<0.05) are in bold. GAMLSS-BEZI: Generalized Additive Models for Location, Scale and Shape (GAMLSS) with a zero inflated beta (BEZI) family; LMAS: linear model with arcsin squareroot transformation (implemented in the software MaAsLin); FDR: false discovery rate.

*Example 3: Effect of diarrheal episodes on bacterial phyla stratified by duration of exclusive breastfeeding (EBF)*

Figure 4 showed average of relative abundance of bacterial phyla in groups of infants from 6 months to 2 years of age with vs. without diarrhea around the time of stool sample collection stratified by duration of EBF. In infants who received less than two months of EBF, a higher abundance of Firmicutes and a lower abundance of Actinobacteria was observed in the groups of infants with diarrhea vs. those without diarrhea (Figure 4, upper panel). GAMLSS-BEZI detected a significant difference in both Firmicutes and Actinobacteria. In contrast, in infants who received more than two months of EBF, no difference in relative abundance of any bacterial phylum was observed between those with diarrhea vs. those without diarrhea (Figure 4, lower panel) and GAMLSS-BEZI did not report any significant difference (Table 5). This example demonstrates that GAMLSS-BEZI detects differential abundances when there is observed difference and does not report difference when there is no observed difference.

**Figure 4.**
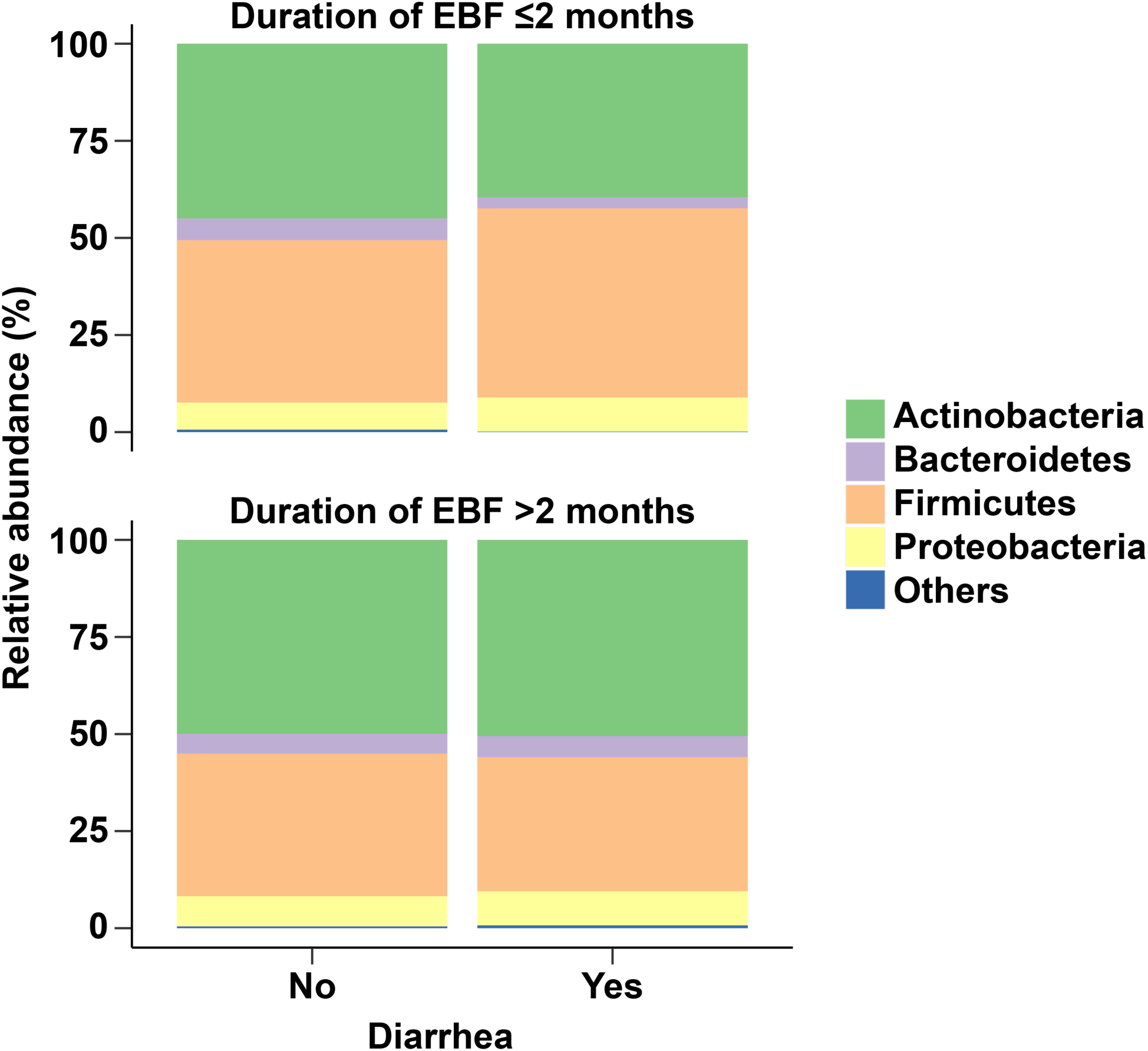
Relative abundance of bacterial phyla in infants from 6 months to 2 years of age with diarrhea vs. without diarrhea at the time of stool sample collection stratified by duration of exclusive breastfeeding (EBF). Data from Bangladesh study.

**Table 5.** Results of GAMLSS-BEZI and LMAS: real microbiome data example 3 (diarrhea vs. no diarrhea stratified by duration of EBF).

| Bacterial phyla | GAMLSS-BEZI |  |  |  |  | LMAS |  |  |  |  |
| --- | --- | --- | --- | --- | --- | --- | --- | --- | --- | --- |
|  | Estimate | 95% Lower limit | 95% Upper limit | p-value | FDR adjusted p-value | Estimate | 95% Lower limit | 95% Upper limit | p-value | FDR adjusted p-value |
| *In infants with duration of exbf <=2 months (diarrhea vs. no diarrhea comparison)* |  |  |  |  |  |  |  |  |  |  |
| Actinobacteria | -0.73 | -1.12 | -0.34 | **0.0003** | 0.0011 | -0.12 | -0.23 | 0.0 | **0.0424** | 0.0848 |
| Bacteroidetes | -0.29 | -0.68 | 0.10 | 0.1524 | 0.2032 | 0.06 | -0.12 | 0.01 | 0.0852 | 0.1136 |
| Firmicutes | 0.49 | 0.15 | 0.84 | **0.0055** | 0.0109 | 0.11 | 0.01 | 0.2 | **0.0269** | 0.0848 |
| Proteobacteria | -0.17 | -0.54 | 0.20 | 0.3729 | 0.3729 | 0.00 | -0.07 | 0.08 | 0.9060 | 0.9060 |
| *In infants with duration of exbf >2 months (diarrhea vs. no diarrhea comparison)* |  |  |  |  |  |  |  |  |  |  |
| Actinobacteria | 0.02 | -0.42 | 0.46 | 0.9243 | 0.9243 | 0.00 | -0.10 | 0.10 | 0.9626 | 0.9989 |
| Bacteroidetes | 0.07 | -0.41 | 0.56 | 0.7680 | 0.9243 | 0.01 | -0.07 | 0.09 | 0.8101 | 0.9707 |
| Firmicutes | -0.02 | -0.40 | 0.36 | 0.9142 | 0.9243 | -0.01 | -0.13 | 0.12 | 0.8927 | 0.9707 |
| Proteobacteria | 0.12 | -0.33 | 0.56 | 0.6043 | 0.9243 | 0.02 | -0.06 | 0.11 | 0.5875 | 0.9191 |

Data from Bangladesh study. Comparison of bacterial relative abundances at phylum level in infants from 6 months to 2 years of age with diarrhea vs. no diarrhea at the time of stool sample collection stratified by duration of exclusive breastfeeding. Significant p-values (<0.05) are in bold. GAMLSS-BEZI: Generalized Additive Models for Location, Scale and Shape (GAMLSS) with a zero inflated beta (BEZI) family; LMAS: linear model with arcsin squareroot transformation (implemented in the software MaAsLin); FDR: false discovery rate.

#### Illustration of meta-analysis workflow with real microbiome data from four studies

We show examples for the comparison of gut microbiomes between male vs. female infants ≤ 6 months of age adjusting for feeding status and infant age at time of sample collection. We illustrate the meta-analyses across microbiome studies with data from four studies (total number of stool samples=610 [female=339, male=271]). These four studies include one in Bangladesh [14], one in Haiti [11], and two in the USA (California and Florida [CA_FL] and North Carolina [NC]) [12, 28] (Table 6, Additional file 1).

**Table 6.**
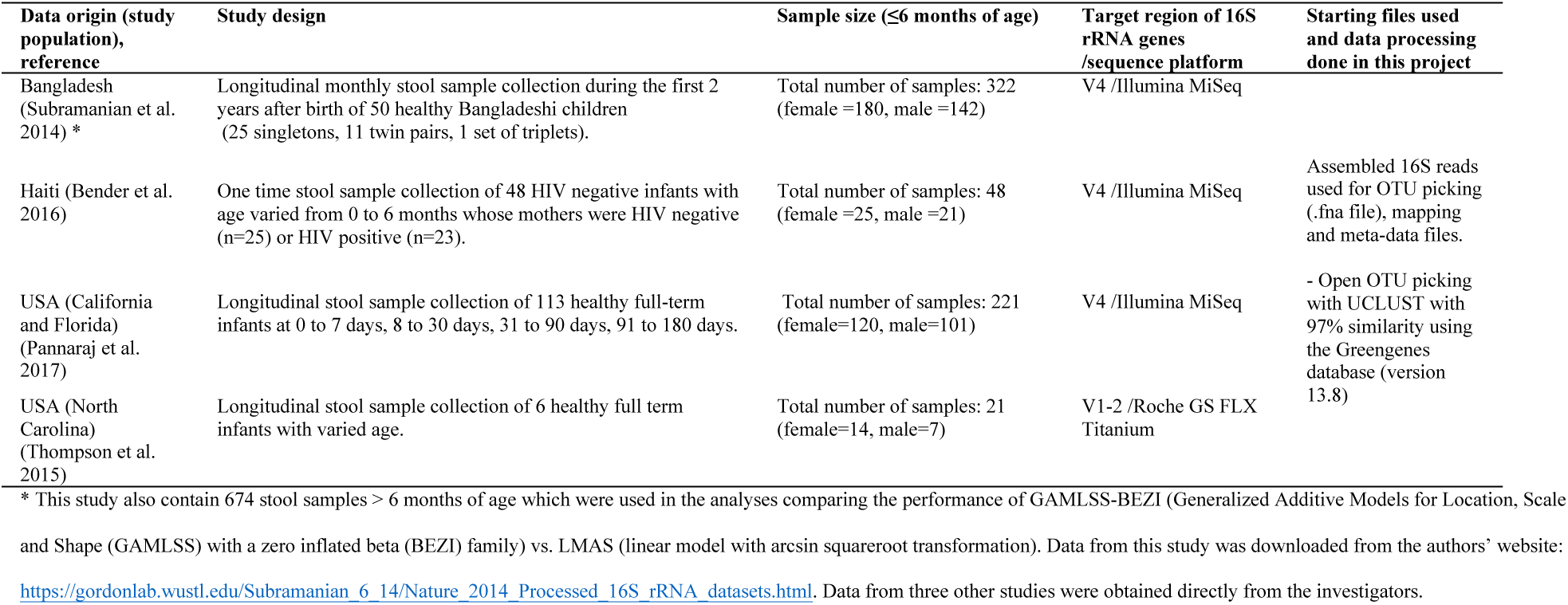
Summary of four studies included in meta-analysis.

#### Alpha diversity

For each study, the *alpha.compare* function imports the outputs from “*alpha_rarefaction.py”* QIIME1 script and calculates mean alpha diversity for different indices for each sample based on a user defined rarefaction depth. Mean alpha diversity indexes are standardized to have a mean of 0 and standard deviation of 1 to make these measures comparable across studies. Standardized alpha diversity indexes are compared between groups adjusting for covariates using LM. Meta analysis across studies is then done and the results are displayed as a standard meta-analysis forest plot (Additional file 1). Our results showed that alpha diversity (four commonly used indexes Shannon, Phylogenetic diversity whole tree, Observed species, Chao1) was not different between male and female infants ≤6 months of age in the meta-analysis of the four included studies (pooled standardized alpha diversity (Shannon index) difference (DD)= −0.01 standard deviation (sd), 95% CI−[−0.15; 0.12], p-value=0.8288) (Figure 5, Additional file 1).

**Figure 5.**
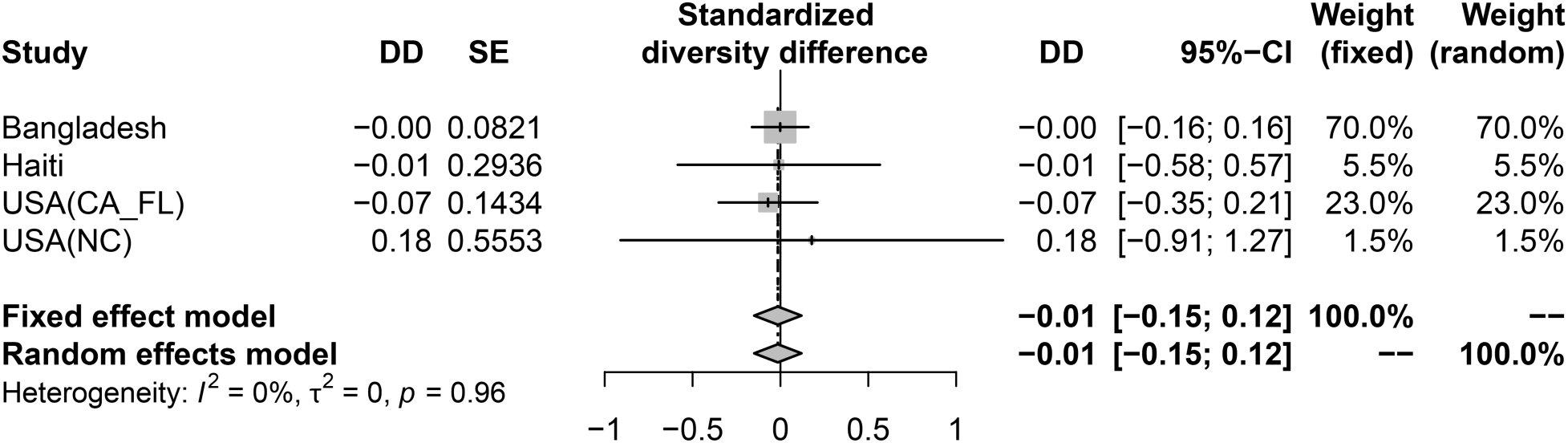
Meta-analysis of four included studies: the difference in gut microbial diversity between male vs. female infants ≤ 6 months. The difference in gut microbial alpha diversity (standardized Shannon index) between male vs. female infants ≤ 6 months old from each study and the pooled effect across four included studies (meta-analysis). Estimates for diversity difference and corresponding standard error from each study were from linear mixed effect models (for longitudinal data) or linear models (for non-longitudinal data) and were adjusted for feeding status and age of infants at sample collection. Pooled estimates of standardized diversity difference and their 95% CI were from random effect meta-analysis models with inverse variance weighting and DerSimonian–Laird estimator for between-study variance based on the adjusted estimates and corresponding standard errors of all included studies (results of fixed effect meta-analysis model are also showed). USA: United States of America; CA: California; FL: Florida; NC: North Carolina; DD: standardized diversity difference; SE: Standard error.

#### Microbiota age

A RF model based on the relative abundance of the shared bacterial genera of the four included studies explained 96% of the variance related to chronological age in the training set and 67% of the variance related to chronological age in the test set of Bangladesh data. This performance is better than the original RF model proposed by Subramanian et al [14]. The predicted infant age in each included study based on relative abundance of the shared gut bacterial genera using this RF model is referred to as gut microbiota age. Standardized gut microbiota age are compared between groups adjusting for covariates using LM and meta-analysis across studies is then done. Our results showed that standardized microbiota age was significantly different between genders but in opposite directions in two studies with small sample sizes (Haiti and North Carolina).

However, meta-analysis of all four studies revealed no significant difference in gut microbiota age between genders after adjusting for feeding status and infant age at time of sample collection (pooled standardized microbiota age difference (MD)=-0.06 sd, 95% CI=[-0.32; 0.20], p-value=0.6348) (Figure 6, Additional file 1).

**Figure 6.**
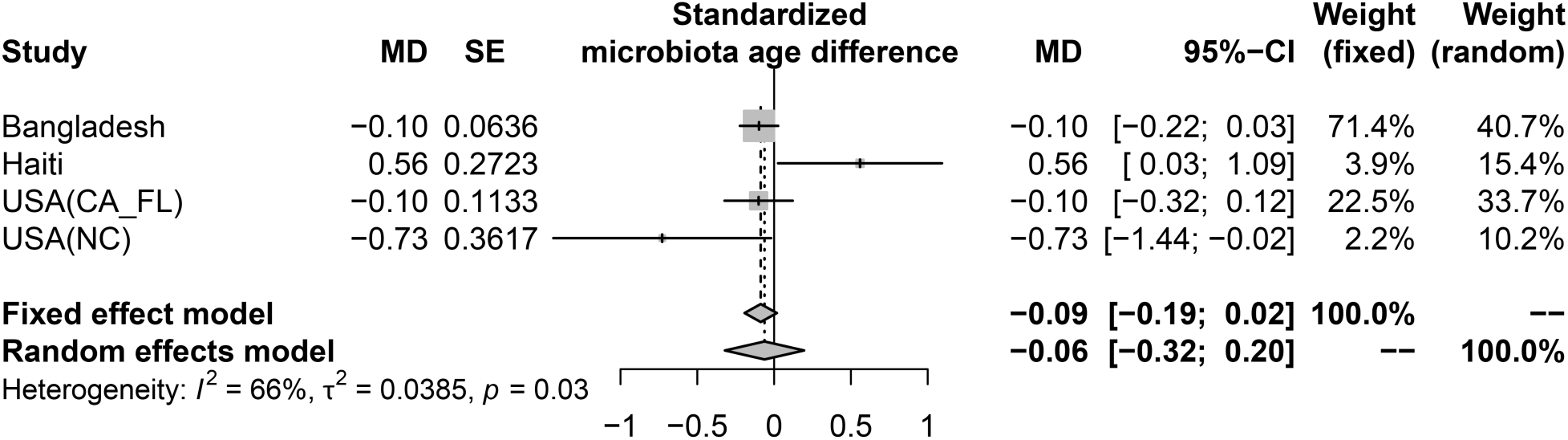
Meta-analysis of four included studies: the difference in gut microbiota age between male vs. female infants ≤ 6 months. The difference in gut (standardized) microbiota age between male vs. female infants ≤ 6 months old from each study and the pooled effect across four included studies (meta-analysis). Estimates for standardized microbiota age difference and corresponding standard error from each study were from linear mixed effect models (for longitudinal data) or linear models (for non-longitudinal data) and were adjusted for feeding status and age of infants at sample collection. Pooled estimates of standardized microbiota age difference and their 95% CI were from random effect meta-analysis models with inverse variance weighting and DerSimonian–Laird estimator for between-study variance based on the adjusted estimates and corresponding standard errors of all included studies (results of fixed effect meta-analysis model are also showed). USA: United States of America; CA: California; FL: Florida; NC: North Carolina; MD: standardized microbiota age difference; SE: Standard error.

#### Bacterial taxa relative abundance

The adjusted estimates (log(odds ratio) of relative abundance between genders) from GAMLSS-BEZI for each bacterial taxon and the pooled adjusted estimates across studies (meta-analysis) are displayed as a heatmap (Figure 7, left panel). The adjacent forest plot (right panel) displays the pooled adjusted estimates and their 95%CI with different colors and shapes to reflect the magnitude of pooled p-values. Our meta-analyses of four included studies showed that, after adjusting for feeding status and infant age at time of sample collection, there was no significant difference in bacterial taxa relative abundance from phylum to genus levels between male vs. female infants ≤6 months of age after adjusting for multiple testing except for genus *Coprococcus* whose relative abundance was significantly higher in male vs. female infants (FDR adjusted pooled p-value<0.0001) (Figure 7, Additional file 1).

**Figure 7.**
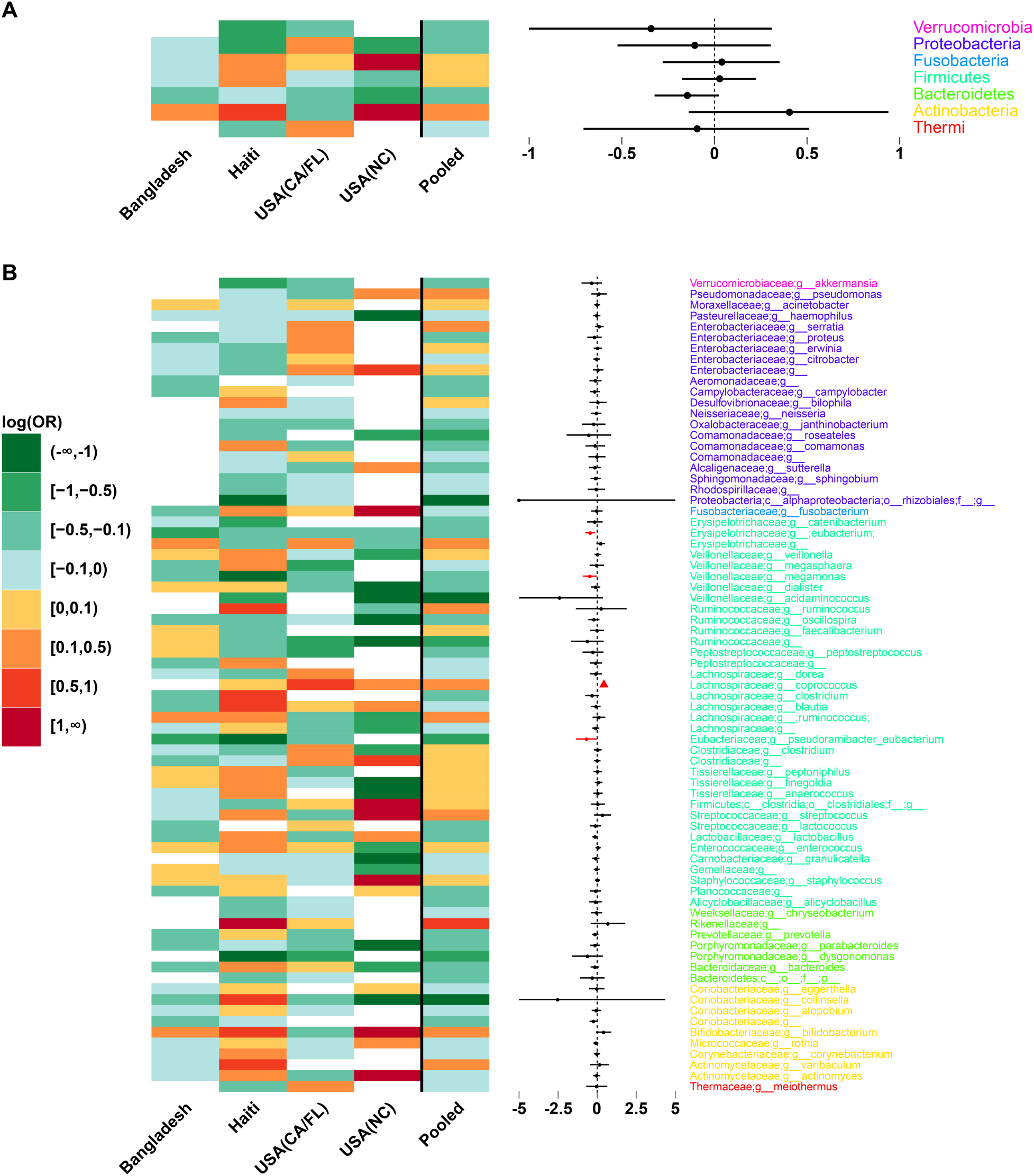
Meta-analysis of four included studies: the difference in relative abundances of gut bacterial taxa between male vs. female infants ≤ 6 months of age. A: Phylum level: heatmap of log(odds ratio) (log(OR)) of relative abundances of all gut bacterial phyla between male vs. female infants for each study and pooled estimates across all studies with 95% confidence intervals (95% CI) (forest plot). B: Genus level: heatmap of log(OR) of relative abundances of all gut bacterial genera between male vs. female infants for each study and pooled estimates across all studies with 95% CI (forest plot). All log(OR) estimates of each bacterial taxa from each study were from Generalized Additive Models for Location Scale and Shape (GAMLSS) with beta zero inflated family (BEZI) and were adjusted for feeding status and age of infants at sample collection. Pooled log(OR) estimates and 95% CI (forest plot) were from random effect meta-analysis models with inverse variance weighting and DerSimonian–Laird estimator for between-study variance based on the adjusted log(OR) estimates and corresponding standard errors of all included studies. Pooled log(OR) estimates with pooled p-values<0.05 are in red and those with false discovery rate (FDR) adjusted pooled p-values <0.1 are in triangle shape. Missing (unavailable) values are in white. USA: United States of America; CA: California; FL: Florida; NC: North Carolina.

#### Bacterial functional (KEGG) pathway relative abundance

Meta-analysis of four included studies showed no difference in relative abundances of bacterial KEGG pathways at level 2 and level 3 between genders after adjusting for feeding status and infant age at time of sample collection and adjusting for multiple testing (Figure 8, Additional file 1).

**Figure 8.**
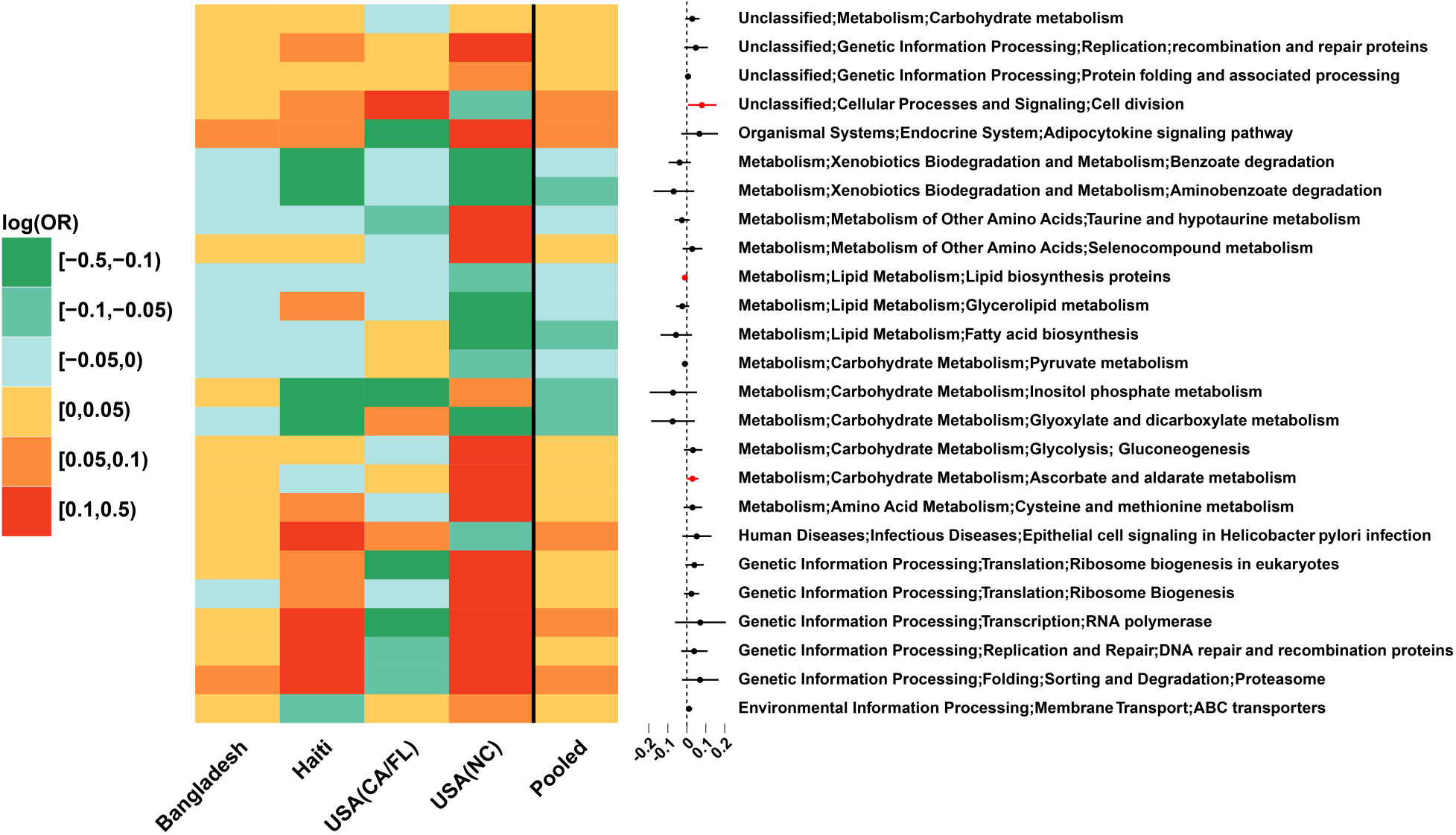
Meta-analysis of four included studies: the difference in relative abundances of gut microbial KEGG (Kyoto Encyclopedia of Genes and Genomes) pathways between male vs. female infants ≤ 6 months of age. Heatmap of log(odds ratio) (log(OR)) of relative abundances of gut microbial KEGG pathways at level 3 between male vs. female infants for each study and pooled estimates of all studies with 95% confidence intervals (forest plot). Only pathways with pooled p-value <0.3 are shown. All log(OR) estimates of each pathway from each study were from Generalized Additive Models for Location Scale and Shape (GAMLSS) with beta zero inflated family (BEZI) and were adjusted for feeding status and age of infants at sample collection. Pooled log(OR) estimates and 95% confidence intervals (95%CI) (forest plot) were from random effect meta-analysis models with inverse variance weighting and DerSimonian–Laird estimator for between-study variance based on the adjusted log(OR) estimates and corresponding standard errors of all included studies. Pooled log(OR) estimates with pooled p-values<0.05 are in red and those with false discovery rate (FDR) adjusted pooled p-values <0.1 were in triangle shape. USA: United States of America; CA: California; FL: Florida; NC: North Carolina.

Difference in gut microbial composition between genders has been reported in adults [29–31] and in some neonatal studies albeit with small sample sizes [32, 33]. However, the reported findings have varied between these studies. Our meta-analyses of four studies showed virtually no difference in gut microbiomes between male vs. female infants ≤6 months of age after adjusting for feeding status and infant age at time of sample collection as well as adjusting for multiple testing. There was one exception: relative abundance of *Coprococcus* was significantly higher in male vs. female infants. *Coprococcus* has been implicated in many conditions including hypertension and autism [34, 35], and the detected difference in our study may provide some insights into the known sex biases in health outcomes.

### CONCLUSION

Our *metamicrobiomeR* package implemented GAMLSS-BEZI for analysis of microbiome relative abundance data and random effect meta-analysis models for meta-analysis across microbiome studies. The advantages of GAMLSS-BEZI are: 1) it directly and properly address the distribution of microbiome relative abundance data which resemble a zero-inflated beta distribution; 2) it has better power to detect differential relative abundances between groups than the commonly used approach LMAS; 3) the estimates from GAMLSS-BEZI are log(odds ratio) of relative abundances between groups and thus are directly comparable across studies.

Moreover, standardization of alpha diversity indexes and predicted microbiota age also yields comparable metrics for cross-study comparisons. Random effects meta-analysis models can be directly applied to pool these comparable adjusted estimates and their standard errors across studies. This approach allows examination of study-specific effects, heterogeneity between studies, and the overall pooled effects across microbiome studies. The examples and workflow using our “*metamicrobiomeR”* package are reproducible and applicable for the analysis and meta-analysis of other microbiome studies.

## AVAILABILITY and REQUIREMENTS

Project name: metamicrobiomeR

Project home page: https://github.com/nhanhocu/metamicrobiomeR

Operating system(s): Platform independent

Programming language: R

Other requirements: R 3.4.2 or higher License: GNU GPL v. 2.

Any restrictions to use by non-academics: none.

## ABBREVIATIONS

BEZI: zero inflated beta
CA: California
CI: Confidence interval
DD: Diversity difference
EBF: exclusive breastfeeding (or exclusively breastfed)
FDR: False Discovery Rate
RF: Random Forest
FL: Florida
GAMLSS: Generalized Additive Models for Location, Scale and Shape
GAMLSS-BEZI: Generalized Additive Models for Location, Scale and Shape with a zero inflated beta family
KEGG: Kyoto Encyclopedia of Genes and Genomes LM: linear/linear mixed effect models
LMAS: linear/linear mixed effect models with arcsin squareroot transformation MD: microbiota age difference
NC: North Carolina
Non-EBF: non exclusive breastfeeding (or non exclusively breastfed)
sd: standard deviation
USA: United States of America

## DECLARATIONS

### Ethics approval and consent to participate

Not applicable

### Consent for publication

Not applicable

### Availability of data and material

All data used in this study are included in these published articles and their supplementary information files (references: [11, 12, 14, 28]). The data from the Bangladesh study [14] were downloaded from the authors’ website: https://gordonlab.wustl.edu/Subramanian_6_14/Nature_2014_Processed_16S_rRNA_datasets.html. The data from three other studies were obtained directly from the investigators. All example datasets, documentations and source code of the “*metamicrobiomeR”* package are available at https://github.com/nhanhocu/metamicrobiomeR.

### Authors’ contribution

NTH conceived the ideas, wrote the R package, documentations, performed the simulations and analyses with inputs from FL.

## Acknowledgements

We would like to thank Dr. Grace M. Aldrovandi (University of California at Los Angeles) and Dr. M. Andrea Azcarate-Peril (University of North Carolina at Chapel Hill) for providing the data used in the examples.

